# An *in vitro* platform with self-sustaining trans-epithelial oxygen gradient to model intestinal barrier function modulation

**DOI:** 10.64898/2026.04.13.718107

**Authors:** Alessandra Maria Anna Rando, Martina Poppa, Chiara Russo, Gianfranco Beniamino Fiore, Monica Soncini

**Affiliations:** Department of Electronics, Information and Bioengineering, Politecnico di Milano, Milan, Italy

**Keywords:** advanced *in vitro* model, oxygen gradient, *in vitro* permeability assays, drug absorption, small intestine, intestinal *in vitro* model

## Abstract

*In vitro* models of the small intestine frequently neglect the role of physiological oxygen tension in drug absorption studies. To meet the throughput demands of preclinical drug development while maintaining compatibility with standard permeability assays, we engineered a new technology, termed Gradient-on-Platform. This system enables the establishment of physiologically relevant oxygen gradients across intestinal epithelium *in vitro* models by leveraging Caco-2 cells metabolic oxygen consumption. In this work, we demonstrated that exposure to physiomimetic oxygen conditions modulates epithelial barrier function, mitigating the excessively tight Caco-2 phenotype typically observed under conventional aerobic cultures, which leads to underestimation of drug bioavailability *in vitro*. Indeed, while complete hypoxia disrupts epithelial barriers, oxygen gradients drive Caco-2 TEER toward values more consistent with *ex vivo* measurements and result in a 3-fold increase in paracellular permeability compared to fully aerobic controls. The incorporation of physiologically relevant oxygen gradients into a scalable, assay-compatible platform could represent a pivotal step toward improving the predictive accuracy of *in vitro* preclinical drug absorption assays.

## 1. Introduction

The intestinal epithelium acts as a selectively permeable barrier, regulating nutrients and molecules absorption, while preventing pathogens from entering the bloodstream. To maximize its absorptive surface area, it is organized into invaginations known as crypts and finger-like protrusions called villi^[1]^. The intestinal tissue is shaped by a finely regulated gaseous microenvironment, in which oxygen plays a central role in tissue homeostasis^[2]^. *In vivo* measurements reveal steep transepithelial oxygen gradients along the crypt–villus axis, with partial pressures decreasing from approximately 85 mmHg in the subepithelial stroma underlying the crypt base to 10-30 mmHg in the small intestinal lumen, and from 42 mmHg to 3-11 mmHg in the colon^[3–5]^. This physiological hypoxia arises from countercurrent oxygen exchange between ascending arterioles and adjacent descending venules along the crypt-villus axis^[6]^, as well as from luminal microbial oxygen consumption^[7]^, and is essential for epithelial metabolism, nutrient absorption^[8]^, barrier integrity maintenance^[9]^, and immune regulation^[10]^. The role played by oxygen availability in modulating transporter activity and epithelial integrity, directly influences intestinal permeability and drug uptake^[11]^, thus making oxygen tension recapitulation a key feature for biomimetic *in vitro* models for drug absorption applications.

However, although oxygen tension has been shown to significantly affect intestinal barrier properties and drug permeability^[12]^, *in vitro* models used for drug absorption studies do not recapitulate physiological oxygen conditions^[13–16]^. Several strategies have nonetheless been developed to control oxygen levels *in vitro*, typically relying on microfluidic platforms incorporating one or more gas channels^[17–19]^. In these systems, defined gas mixtures flow adjacent to liquid culture channels, and local oxygenation conditions are imposed thanks to diffusion through gas-permeable materials, such as polydimethylsiloxane^[17–24]^.

While capable of generating controlled oxygen gradients, such approaches are resource-intensive and technically complex^[25]^. They require bulky auxiliary equipment, including pumps, tubing, valves, and control units, or dedicated hypoxia chambers^[26]^. Moreover, these systems are prone to operational instability, as gas supplies must be continuously monitored and replaced, gas connections are susceptible to leakage, equilibration times are long, and oxygen levels are difficult to maintain consistently over the duration of experiments, thus limiting the large-scale adoption of oxygen gradient-based *in vitro* platforms^[27]^.

Alternative strategies to establish oxygen gradients *in vitro* have therefore focused on simpler, self-sustained approaches that exploit cellular respiration, avoiding the need for external gas perfusion, medium deoxygenation, or anoxic conditions handling. For example, Chen et al.^[28]^ developed a 3D human intestinal model based on a silk protein scaffold with a hollow lumen, in which the interplay between oxygen diffusion and cellular metabolic activity enabled the autonomous formation of an *in vivo*-like oxygen tension gradient along the lumen. Similarly, Kim et al.^[29]^ applied this principle by engineering a bicompartmental culture device incorporating an oxygen-impermeable luminal plug to induce apical deoxygenation. Notably, this design closely resembles standard culture inserts, thereby facilitating assay parallelization and protocol translatability. Microfluidic chip oxygenation can also be limited by coating device walls with oxygen-impermeable materials, relying on cellular consumption for medium deoxygenation^[30]^. Intermediate approaches have also been reported^[31]^, combining diffusion-limited device architectures with medium pre-equilibrated under anaerobic conditions to support gradient formation. However, to the best of our knowledge, self-sustaining oxygen gradients strategies remain largely unexplored in *in vitro* permeability studies.

In this work, we aimed at designing *in vitro* models able to self-sustain oxygen gradients without requiring specific lab equipment, while providing high experimental throughput, specifically for *in vitro* permeability studies applications. Hence, we engineered a new technology, termed Gradient-on-Platform (GoP), to recreate physiological intestinal hypoxia within compartmentalized *in vitro* models by leveraging metabolic oxygen consumption through cellular respiration. The deoxygenation system was coupled with bicompartmental and tricompartmental inserts, namely TTOP^[32]^ and Tri-TOP^[33]^ inserts, respectively, to allow for tissue-specific oxygen tension regulation for permeability testing purposes. The TTOP device incorporates a polycarbonate (PC) microporous membrane that supports cell growth and defines distinct apical and basolateral compartments. On the other hand, the Tri-TOP platform features two PC membranes to generate three physically separated compartments, introducing an intermediate dedicated stromal compartment, which in this configuration is typically modeled using Matrigel.

The GoP’s capability to achieve tissue-relevant oxygen concentrations was evaluated through computational simulations and experimentally validated by using oxygen microsensors. This platform was specifically engineered to optimize absorption studies feasibility by maintaining the design and operating procedures comparable to standard bicompartmental inserts^[34]^. Paracellular transport studies were performed, to evaluate the suitability of the new technology to recapitulate the crucial role played by oxygen tension in modulating Caco-2 barrier function^[9,35–37]^.

## 2. Methods

### 2.1. Gradient-on-Platform device design and fabrication

The Gradient-on-Platform design aimed at enabling controlled hypoxia within the luminal compartment of intestinal epithelium models through an apical deoxygenation module, while providing higher oxygenation in the basolateral compartment, thereby mimicking small intestine physiological conditions. The deoxygenation was driven by metabolic oxygen consumption of intestinal epithelial cells cultured on apical PC membranes of tricompartmental devices (Tri-TOP), beneath which a 575 µm-thick Matrigel layer was introduced to model the stromal component. The apical deoxygenation module consisted of two primary components: a threaded sealing plug and a threaded apical well (Figure 1a). The sealing plug integrated an O-ring (Figure 1a, upper portion), while the apical well incorporated a triangular groove to optimize O-ring compression and achieve a robust oxygen-impermeable seal (Figure 1a, lower portion). To improve plug’s handling and facilitate the assembly of the system, the sealing plug was designed with peripheral grooves, allowing for easier manipulation with tweezers and secure tightening of the components.

**Figure 1.**
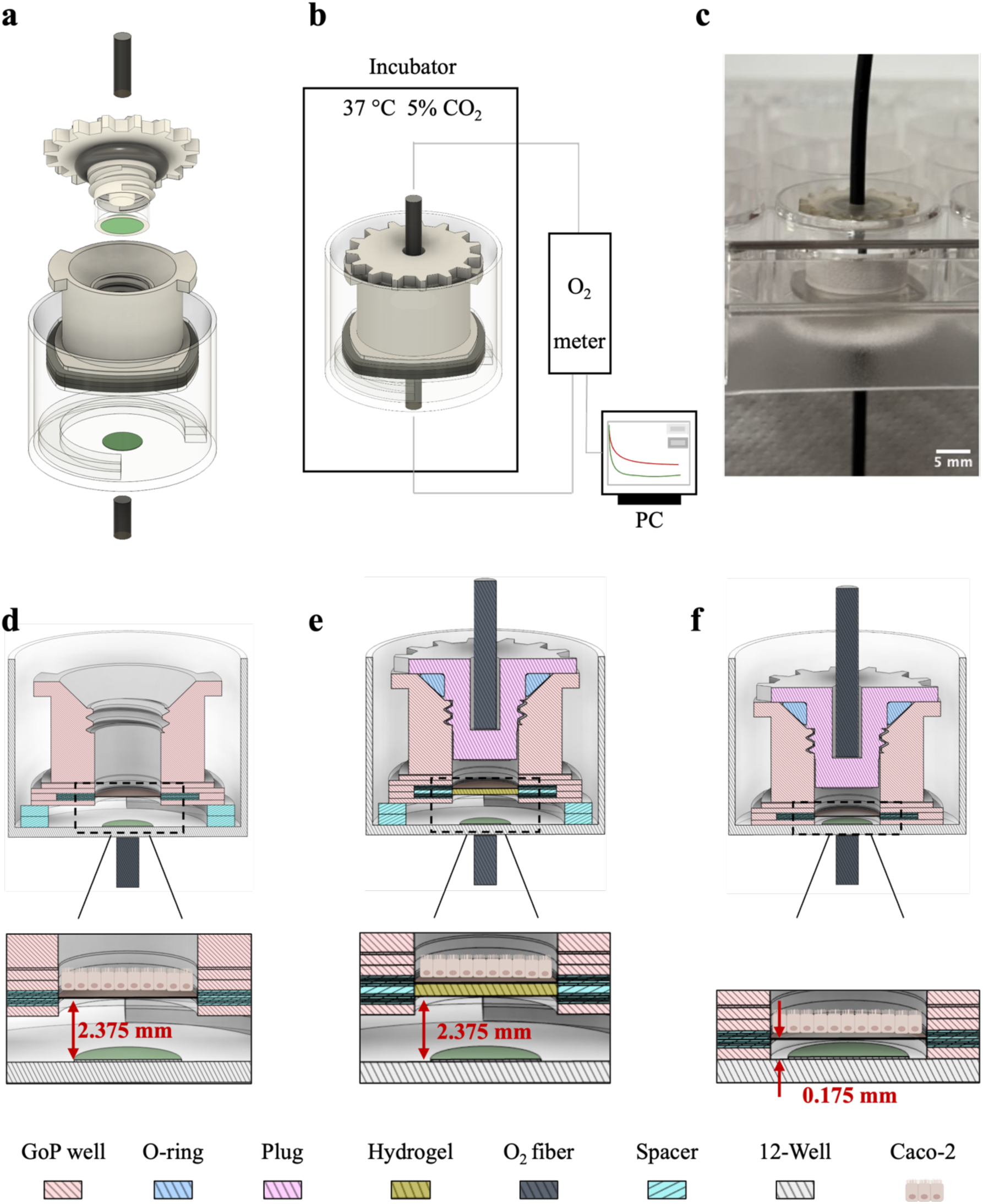
GoP design and oxygen monitoring. (a) CAD model of the GoP system hosted in a well of the 12-well plate integrating oxygen sensors and fibers (green circles represent oxygen sensor spots, black cylinders represent oxygen optical fibers); (b) Schematic representation of oxygen concentration profiles acquisition system; (c) GoP prototype within a 12-well plate with apical and basal oxygen optical fibers. Scale bar = 5 mm; (d) CAD xz-section of hyperoxic, (e) physioxic and (f) hypoxic designs. The zoom-ins highlight Caco-2 cells and hydrogel positioning in the *in vitro* models with respect to the 12-well bottom.

The threaded apical well was 3D printed in polylactic acid (PLA) using an Ultimaker 3 3D printer (Ultimaker B.V., The Netherlands). The sealing plug was fabricated using a Stratasys Objet30 Pro 3D printer (Stratasys, Israel) with VeroClear photopolymer resin, except for its lower portion which consisted of a CO_2_ laser cut (VLS3.50, Scottsdale, USA) completely transparent PMMA (Evonik Röhm GmbH, Germany) cylinder, 6.3 mm in diameter, where a 5 mm oxygen sensor spot was adhered (Figure 1a – upper portion), to enable signal readability via an oxygen optical fiber. Although the apical deoxygenation module was engineered to reduce oxygen tension in the apical luminal compartment, the generation of an oxygen gradient required coordinated basal oxygen supply. To enhance basal oxygen availability, 2.2 mm PMMA C-shaped spacers were adhered at the bottom of the 12-well plate wells, increasing the basal medium volume (Figure 1e) and enhancing diffusional oxygen transport from the lateral compartment. To establish oxygen gradients, the deoxygenation module was integrated with the Tri-TOP platform, which incorporated a 575 µm-thick hydrogel beneath the epithelial monolayer, mimicking the stromal compartment. This hydrogel, specifically Matrigel (Corning, USA), exhibiting a lower oxygen diffusion coefficient (D) than the water-based culture medium^[38]^, introduced an additional diffusional resistance that modulated oxygen flux from the basal compartment.

In addition, two control levels of oxygenation were implemented in the study, by including hyperoxic and deeply hypoxic conditions in two purpose-built auxiliary devices. Hyperoxic samples were obtained by using bicompartmental open-well TTOPs, positioned over the 2.2 mm basal spacers to maximize samples oxygenation (Figure 1d), thereby keeping the oxygen level in the whole system as close as possible to that of the incubator atmosphere. In turn, hypoxic samples were obtained by coupling the apical deoxygenation module with the bicompartmental TTOP, without employing any additional diffusional resistance, and avoiding any spacer in the basolateral compartment to minimize basal medium volume (distance of the membrane from the well bottom of 0.175 mm) (Figure 1f); this configuration was devised to extend deoxygenation to all the inner volume of the device, both apically and basally.

For experimental oxygen concentration characterization, the possibility to incorporate oxygen microsensors within the system was considered pivotal, hence the sealing plug integrated a 5 mm diameter PS-Pst7-NAU oxygen sensor spot (PreSens Precision Sensing GmbH, Germany), and featured an hollow cavity intended to host an oxygen fiber to allow for oxygen concentration monitoring of the apical compartment. For the basal compartment, an oxygen sensor spot was adhered at the bottom of the 12-well plate well, and the oxygen optical fiber tip was positioned below the well plate (Figure 1).

### 2.2. Computational simulations of oxygen concentration profiles

To inform GoP geometric design, the temporal evolution of oxygen concentration within the device in the presence of cells was modeled and simulated using COMSOL Multiphysics v.6.4 (COMSOL Inc., USA). Time-dependent analyses of the oxygen concentration profiles were performed to predict steady-state oxygen concentration in the apical and basolateral compartment of the *in vitro* model, characterizing the optimal design features for the recapitulation of oxygen concentrations relevant to the intestinal tissue.

Oxygen transport was modelled as previously described^[39]^ by using the general mass transport equation (Equation 1) where *c* is the oxygen spatial concentration [mol m^-3^], *D* the diffusion coefficient [m² s^-1^], *V* the volumetric consumption rate [mol m^-3^ s^-1^], *P* the volumetric production rate [mol m^-3^ s^-1^], and *v* the fluid velocity [m s^-1^]. The equation was opportunely simplified, according to device’s foreseen operating conditions, under the assumptions of negligible convection in static culture conditions, and negligible production term.

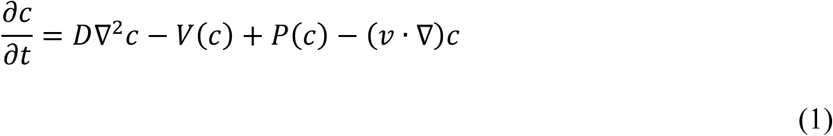

Oxygen consumption by Caco-2 cells was assumed to follow Michaelis-Menten kinetics^[40]^ described by Equation 2, where *V_max_* is the maximum oxygen consumption rate [mol m^-3^ s^-1^], *c* is the spatial oxygen concentration [mol m^-3^], *k_m_* is the Michaelis-Menten constant [mol m^-3^], corresponding to the oxygen concentration where consumption drops to 50% of its maximum.

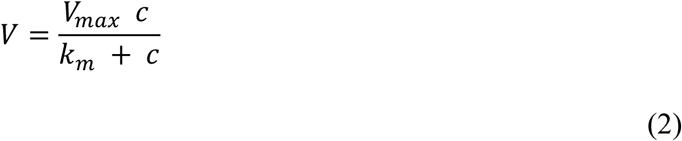

Moreover, a smoothed Heaviside step-down function (δ) was applied to account for cessation of consumption below a critical oxygen concentration (C_cr_).

The GoP computational model was composed of geometrical domains representing the PC microporous membranes hosted in the Tri-TOP platform, the interposed hydrogel, the cellular monolayer positioned on the upper PC microporous membrane, the apical medium above the cell monolayer, the threaded GoP well and an additional domain mimicking the basal medium (Figure 2a and Supplementary Information, Section3, Figure S2).

**Figure 2.**
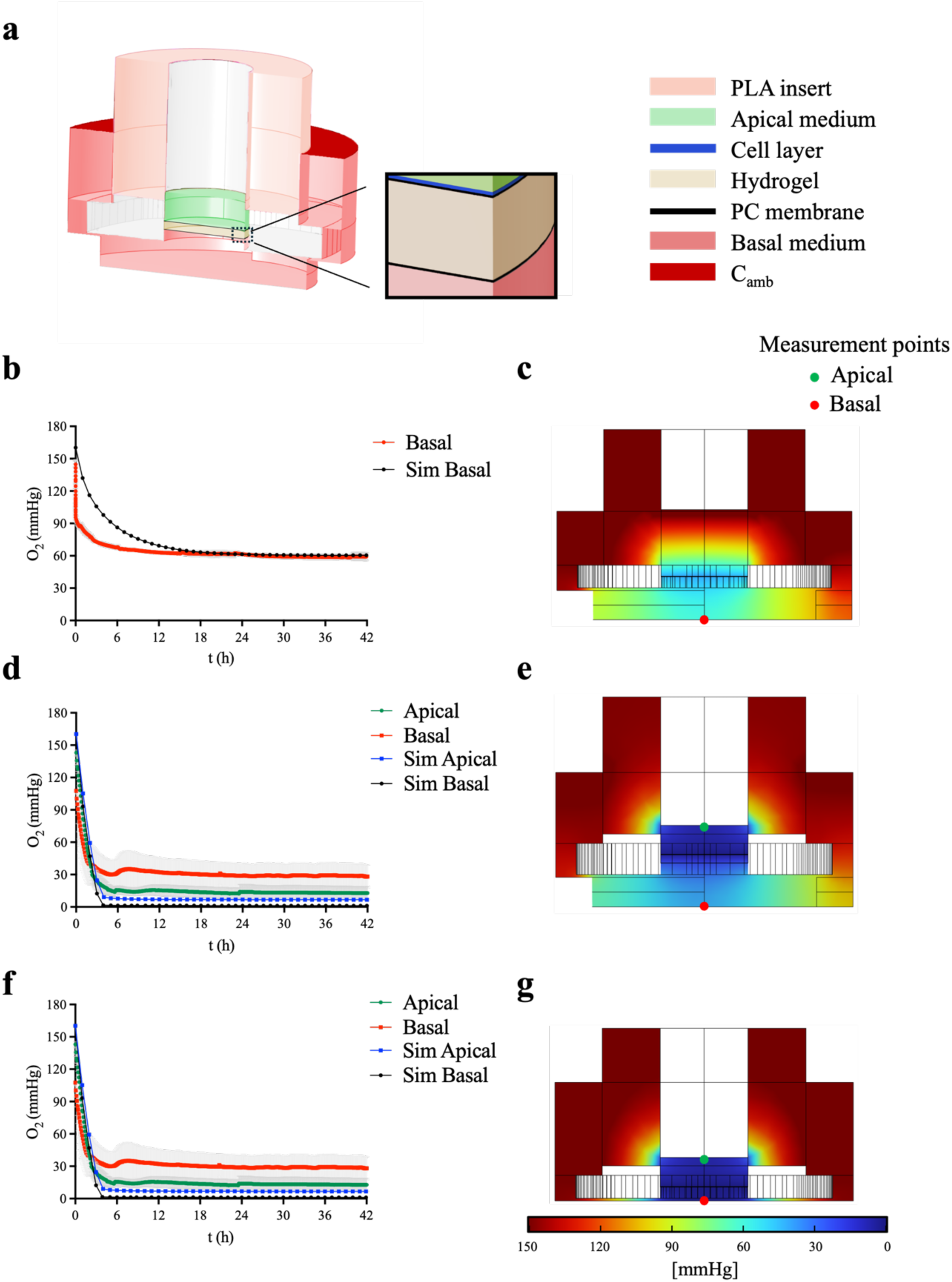
Experimental and computational assessment of intestinal tissue-relevant oxygen concentrations. (a) Schematic of the computational domains used to model the Gradient-on-Platform in COMSOL, with the insert highlighting membranes and cellular domains; (b-g) Oxygen concentration profiles and xz– cross section of simulated oxygen concentration after 42 hours for hyperoxic (b,c), physioxic (d,e) and hypoxic (f,g) conditions. The dotted lines represent simulated data. Experimental data are presented as mean ± SEM (n=3 devices per conditions)

Oxygen diffusion coefficients were assigned to each domain based on material properties (Table 1), assuming purely diffusive transport. The PLA threaded well was considered marginally permeable to oxygen with an oxygen diffusion coefficient experimentally derived (D_PLA_) (Supplementary Information, Section1, Figure S1). The oxygen diffusion coefficient for the PC microporous membranes (D_membrane_) was estimated from the diffusion coefficient of oxygen in bulk polycarbonate, corrected for membrane porosity (Supplementary Information, Section2), while D_Medium_ and D_Matrigel_ were obtained from previously published works. Oxygen consumption was imposed only in the cellular layer and modelled as a product of maximum oxygen consumption rate per cell (R_max_) and total cells in culture area (N), estimated experimentally from nuclei staining.

**Table 1.** Model parameters of the computational simulations.

| Parameter | Value | Units | Description |
| --- | --- | --- | --- |
| $C_0(0)$ | 0.218 | $\text{mol m}^{-3}$ | Initial oxygen concentration<sup>[41]</sup> |
| $C_{\text{amb}}$ | 0.218 | $\text{mol m}^{-3}$ | Atmospheric oxygen concentration<sup>[41]</sup> |
| $D_{\text{Medium}}$ | $3.00 \times 10^{-9}$ | $\text{m}^2 \text{s}^{-1}$ | O<sub>2</sub> diffusion in culture medium<sup>[39]</sup> |
| $D_{\text{Matrigel}}$ | $1.50 \times 10^{-9}$ | $\text{m}^2 \text{s}^{-1}$ | O<sub>2</sub> diffusion in Matrigel<sup>[38]</sup> |
| $D_{\text{Membrane}}$ | $3.60 \times 10^{-10}$ | $\text{m}^2 \text{s}^{-1}$ | O<sub>2</sub> diffusion in microporous PC membranes |
| $D_{\text{PLA}}$ | $2.77 \times 10^{-10}$ | $\text{m}^2 \text{s}^{-1}$ | O<sub>2</sub> diffusion in the PLA threaded well |
| $R_{\text{max}}$ | $2.20 \times 10^{-17}$ | $\text{mol cell}^{-1} \text{s}^{-1}$ | Maximum O<sub>2</sub> consumption rate per cell<sup>[42]</sup> |
| $K_m$ | 0.001 | $\text{mol m}^{-3}$ | Michaelis-Menten constant<sup>[39]</sup> |
| $N$ | Average cell number | cells | Total cells in culture area |
| $V_{\text{cell}}$ | $6.64 \times 10^{-10}$ | $\text{m}^3$ | Volume of the cells |
| $C_{\text{cr}}$ | 0.00005 | $\text{mol m}^{-3}$ | Critical oxygen concentration<sup>[39]</sup> |

An initial oxygen concentration equal to the concentration of oxygen in a water-based medium in equilibrium with oxygen atmospheric concentration was assumed throughout the domain, following Henry’s law^[39]^, reflecting aerobic culture prior to the sealing of the luminal plug (C_0_(0)). This O_2_ concentration was also imposed at the medium surface in contact with air via a constant concentration boundary condition on the top of the domain (C_amb_). The TTOP and Tri-TOP devices, the sealing plug, and the polystyrene 12-well plate were considered oxygen-impermeable, with a no-flux boundary condition.

COMSOL simulations of hyperoxic and deeply hypoxic control conditions were based on the same model, accounting for condition-specific geometrical features (e.g., no spacer in hyperoxic settings) (Supplementary Information, Section3, Figure S2).

### 2.3. Experimental oxygen concentration characterization

Caco-2 cells were maintained in humidified incubator (37°C, 5% CO_2_) in low-glucose Dulbecco’s Modified Eagle Medium (DMEM, Gibco, USA), supplemented with 20% Fetal Bovine Serum (FBS, Euroclone, Italy), 1% MEM Non-Essential Amino Acids (NEAA, Euroclone, Italy) and 1% Penicillin-Streptomycin-Amphotericin B (Euroclone, Italy), and passaged when 70% confluent.

For oxygen concentration characterization experiments, apical PC membranes of bicompartmental TTOP and Tri-TOP inserts were coated with 10 μg ml^-1^ Collagen I (Corning, USA) in 1X Dulbecco’s Phosphate Buffered Saline (1XDPBS, Euroclone, Italy), incubated for 2 hours in the humidified incubator. Tri-TOP stromal compartments were filled with 35μl of 1 mg ml^-1^ Matrigel (Corning, USA) in 1XDPBS and polymerized for 30-60 minutes in the humidified incubator, then 5 × 10^4^ Caco-2 cells were seeded on the collagen-coated PC membrane of each culture inserts, suspended in 200 μl culture medium, while 1.5 ml culture medium was added to the basolateral compartment. Caco-2 cells were allowed to grow until confluence in aerobic (i.e., hyperoxic) conditions.

Oxygen concentration profiles were acquired overtime up to 42 hours^[43]^, while the devices were hosted in the humidified incubator to maintain optimal cell culture conditions. While for physioxic and hypoxic samples both apical and basolateral oxygen concentration profiles were measured (Figure 1e,f), the sealing plug was not employed for hyperoxic controls and continuous oxygenation from the luminal environment was assumed, thus only the basolateral oxygen profile was recorded (Figure1d).

The oxygen optical fibers were connected to the OXY-4 ST (G2) oxygen meter (PreSens Precision Sensing GmbH, Germany) and data were acquired through the PreSens measurement studio 2 software (PreSens Precision Sensing GmbH, Germany) (Figure 1b). Oxygen concentration profiles of three independent devices were then analyzed through GraphPad Prism 10 software.

Following oxygen concentration profiles characterization, samples were fixed with 10% Formalin (Sigma-Aldrich, USA), permeabilized with 0.1% Triton-X (Euroclone, Italy) in 1XDPBS, and stained with and 1:5000 DAPI (Biochemica, Italy). Samples were mounted on microscope slides with 15% glycerol (Sigma-Aldrich, USA) in 1XDPBS, then imaged with the Olympus ix83 wide field fluorescence microscope (Olympus, Japan). Cell density was quantified from DAPI staining by using Fiji ImageJ (National Institutes of Health) software (n=3 samples per condition, n=10 images per sample). The average total number of cells for each condition was modelled in the computational simulations.

### 2.4 Caco-2 barrier tightness modulation

Caco-2 cells were cultured under aerobic conditions until confluence, as previously described. Baseline TEER was measured prior to oxygen modulation to confirm barrier integrity. Oxygen relevant conditions were then established by selectively modifying the system configuration: sealing the apical plug generated physioxic conditions; sealing the apical plug combined with removal of the basal spacer promoted hypoxic conditions by limiting both apical and basal oxygen availability; whereas hyperoxic controls were maintained under aerobic conditions throughout the experiment. Oxygen-relevant conditions were maintained for 48 hours, after which TEER was re-assessed to quantify oxygen-induced alteration of barrier integrity. Samples were then fixed with 10% Formalin, permeabilized with 0.1% Triton-X in 1XDPBS, blocked in 1% BSA in 1XDPBS and incubated with 1:300 anti-ZO1 monoclonal antibody (Invitrogen, USA) in 1% Bovine Serum Albumin (BSA) overnight at 4°C, then 1:100 488 secondary antibody (Invitrogen, USA) and 1:5000 DAPI (Biochemica, Italy) were incubated for 1 hour at room temperature. Samples were mounted on microscope slides with 15% glycerol in 1XDPBS and imaged via Olympus ix83 wide field fluorescence microscope. At least 5 images per sample (n=3 samples per condition) were acquired and analyzed with Fiji ImageJ software for cell density and tight junction formation. Junctional length was quantified by measuring at least 5 distinct tight junction segments per image, as an indicator of monolayer packing density. Shorter junctional lengths were interpreted as reflecting a more compact and tightly packed epithelial organization, whereas longer junctional lengths indicated increased intercellular spacing and reduced packing^[44]^.

### 2.5 Paracellular permeability assay

Paracellular permeability assays were performed employing tetramethylrhodamine (TMR) labeled 40 kDa Dextran. 0.1 mg ml^-1^ TMR-40 kDa dextran stock solution was prepared in transport buffer (TB) consisting of phenol-red free low glucose DMEM (Euroclone, Italy) supplemented with 20% FBS, 1% NEAA, 1% L-glutamine and 1% Penicillin-Streptomycin-Amphotericin B. For the assays, the culture medium was replaced by TB and samples were allowed to equilibrate in TB for 1 hour prior to perform TEER measurements to assess starting monolayer integrity. Then, apical TB was replaced by 200 μl of fluorescent molecule, from which 100 μl samples were immediately taken to assess starting molecule concentration (C_0_). Oxygen relevant conditions were then induced as previously described and samples were transferred to a humidified incubator (37°C, 5% CO_2_). 100 μl samples were then taken from the receiver (basal) compartments at 24 hours and 48 hours. Samples fluorescence was then quantified by a spectrofluorometer (Tecan Spark plate reader, SparkControl Software) with excitation wavelength of 530/20 and emission wavelength of 580/20. TMR-40 kDa dextran transport was quantified against standard dilution curves of known concentrations. The apparent permeability *P_app_* was calculated according to Equation 3, where *dQ/dt* is the transport rate across the cell monolayer, *A* is the surface area and *C_0_* is the starting concentration of the molecule in the donor compartment. TEER was also used as an endpoint to assess barrier functionality at the end of the experiment.

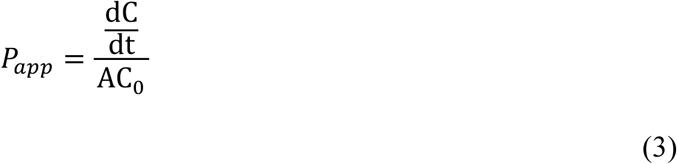

### 2.6 Statistical analysis

Statistical analysis was performed using GraphPad Prism 10 software. Dataset normality was assessed via Shapiro-Wilk test. Sample size (n) and statistical tests are reported in each figure caption. Statistical significance was considered when p<0.05.

## 3. Results

### 2.1. The Gradient-on-Platform module supports the establishment of intestinal physiological oxygen gradients

The GoP module was engineered to reproduce physiological oxygen gradients across intestinal epithelial cell cultures. The target oxygen concentrations for the two compartments were set to ≈1% (≈ 8 mmHg) in the apical compartment, and ≈6-12% (≈ 45-90 mmHg) in the basolateral compartment, according to reference *in vivo* data^[3–5]^.

The design of the GoP module was supported by computational simulations aiming at predicting steady-state oxygen concentration in apical and basal compartments, and experimental oxygen concentration measurements overtime, to assess the effective Caco-2 cells capability of establishing intestinal tissue-relevant oxygen concentrations in the considered setup.

The experimental characterization of the *in vitro* platform showed good agreement between steady-state oxygen tensions in both compartments of physioxic samples with physiological benchmark values^[3–5]^ (Figure 2d). At the same time, confirmed the establishment of negative and positive controls, being hyperoxic (Figure 2b) and hypoxic samples (Figure 2f), respectively.

The computational predictions showed realistic qualitative time courses, reaching steady states in durations on the order of hours, which are comparable with those observed experimentally. Quantitatively, the plateau levels that were reached after each transient were well aligned with experimental data for basal hyperoxic (Figure 2b) and apical physioxic and hypoxic oxygen concentrations (Figure 2d,f). However, experimental measurements of basal oxygen tension in physioxic and hypoxic conditions showed higher oxygenation in the basal compartment with respect to the simulated values. This indicates that, in the scenarios where we introduced intentional resistances to diffusion towards the basal compartment, the hindrance to oxygen transport was less efficient in reality than in the numerical prediction. Nonetheless, experimental data for the physioxic condition still aligned with the chosen target for basal oxygen tension (6-12%^[3–5]^), thus device design was still considered adequate for the foreseen application. Along similar lines, in hypoxic control samples both compartments at steady-state reached oxygen concentrations below 5% (≈ 38 mmHg), which is widely accepted as a hypoxia threshold in cellular and tissue studies^[45–47]^.

### 3.3 Oxygen tension regulates cell organization

Caco-2 cells were cultured under aerobic conditions until confluence and tissue maturation, following established protocols^[48–51]^. Monolayer integrity and tight junction formation were assessed by TEER measurements prior to the induction of oxygen-relevant conditions (Figure 3a, left panel), confirming adequate maturation of the epithelial constructs. The coefficient of variation calculated across the mean TEER values of the three experimental groups (hypoxia, physioxia, and hyperoxia) was 11.6%, indicating low-to-moderate variability and supporting the assumption of comparable starting conditions among samples. Physioxic and hypoxic environments were subsequently applied for 48 hours, while hyperoxic samples were maintained under standard aerobic culture conditions. Exposure to altered oxygen tensions resulted in a pronounced decrease in TEER values (Figure 3a, right panel) and cell density (Figure 3b), accompanied by an increase in tight junction length, indicative of reduced epithelial packing (Figure 3c). Consistently, immunofluorescence imaging revealed a denser and more compact monolayer under hyperoxic conditions (Figure 3d, left panel), which appeared progressively looser under physioxic (Figure 3d, central panel) and, more markedly, hypoxic conditions (Figure 3d, right panel). It is well established that conventional aerobic Caco-2 cultures tend to overestimate the physiological tightness of the intestinal epithelium^[52,53]^, whereas physiological TEER values for the intestinal tissue are typically reported around 100 Ωcm^2^ in *ex vivo* studies^[54,55]^, coherently with what reported here for samples subjected to O_2_ altered regimes (Figure 3a – right panel). Conversely, an excessive reduction in monolayer tightness may be indicative of an inflammatory-like state^[36,56]^, potentially compromising epithelial barrier function. Indeed, TEER decrease below 50% has been previously reported in response to pro-inflammatory stimuli^[32,52,56]^.

**Figure 3.**
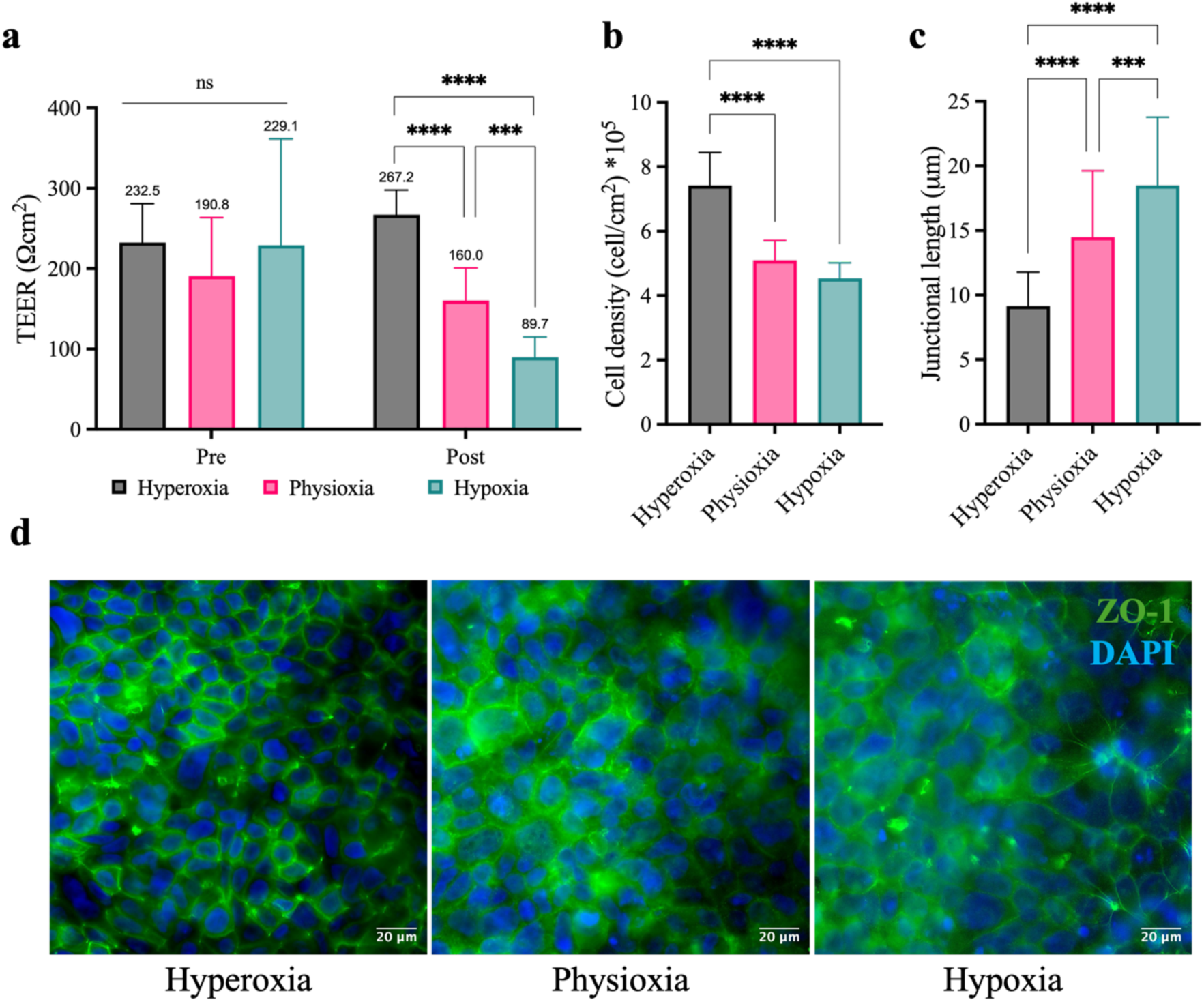
Oxygen-dependent Caco-2 cells monolayer organization. Quantitative assessment of (a) TEER (n=3 samples per condition, 3 measurements per sample), (b) cell density (n=3 samples per condition, at least 5 images per sample), and (c) tight junction length (n=3 samples per condition, at least five images per sample, 5 tight junction segment measurements per image) in different oxygen tension regimes – hyperoxia (grey), physioxia (pink) and hypoxia (teal); (d) Representative images of Caco-2 cells in the considered settings (nuclei in blue (DAPI), ZO-1 in green). Scale bar = 20 μm. Data are presented as mean ± SD. Data normality: Shapiro-Wilk test. Statistics: t-test with Welch’s correction between each pair of conditions. ** p<0.01, *** p<0.001, ****p<0.0001

### 3.4 Physiologically relevant oxygen concentrations modulate Caco-2 barrier function

Barrier integrity assays involving TMR-40 kDa dextran aimed at further investigating oxygen induced modulation of Caco-2 barrier function, by measuring TEER before and after the assay and monitoring paracellular transport while oxygen-relevant conditions were applied (Figure 3a). As previously introduced, oxygen tension emerged as a key determinant of Caco-2 barrier function and paracellular permeability^[35]^. Hyperoxic conditions (grey bar) induced TEER increase after 48 hours (Figure 4b), coupled with low apparent permeability to TMR–40 kDa dextran (Figure 4c), confirming the well-known non-physiological overtightening of Caco-2 monolayers^[48]^. In contrast, exposure to physioxic oxygen (pink bars) levels effectively attenuated this overtight response, restoring TEER and P_app_ values toward a more physiologically relevant permeability profile (Figure 4b,c)^[54]^, with TEER better aligned with physiological benchmark values (119 ± 51 Ωcm^2^)^[54,55]^ and a 3-fold increase in TMR-40 kDa Dextran P_app_. Conversely, hypoxic conditions (teal bars, Figure 4b,c) led to a pronounced loss of barrier integrity, as evidenced by a marked TEER decrease (around 70%) and a concomitant 6-fold increase in P_app_, indicating barrier disruption rather than a controlled modulation of epithelial tightness^[9,36,37,52,56]^.

**Figure 4.**
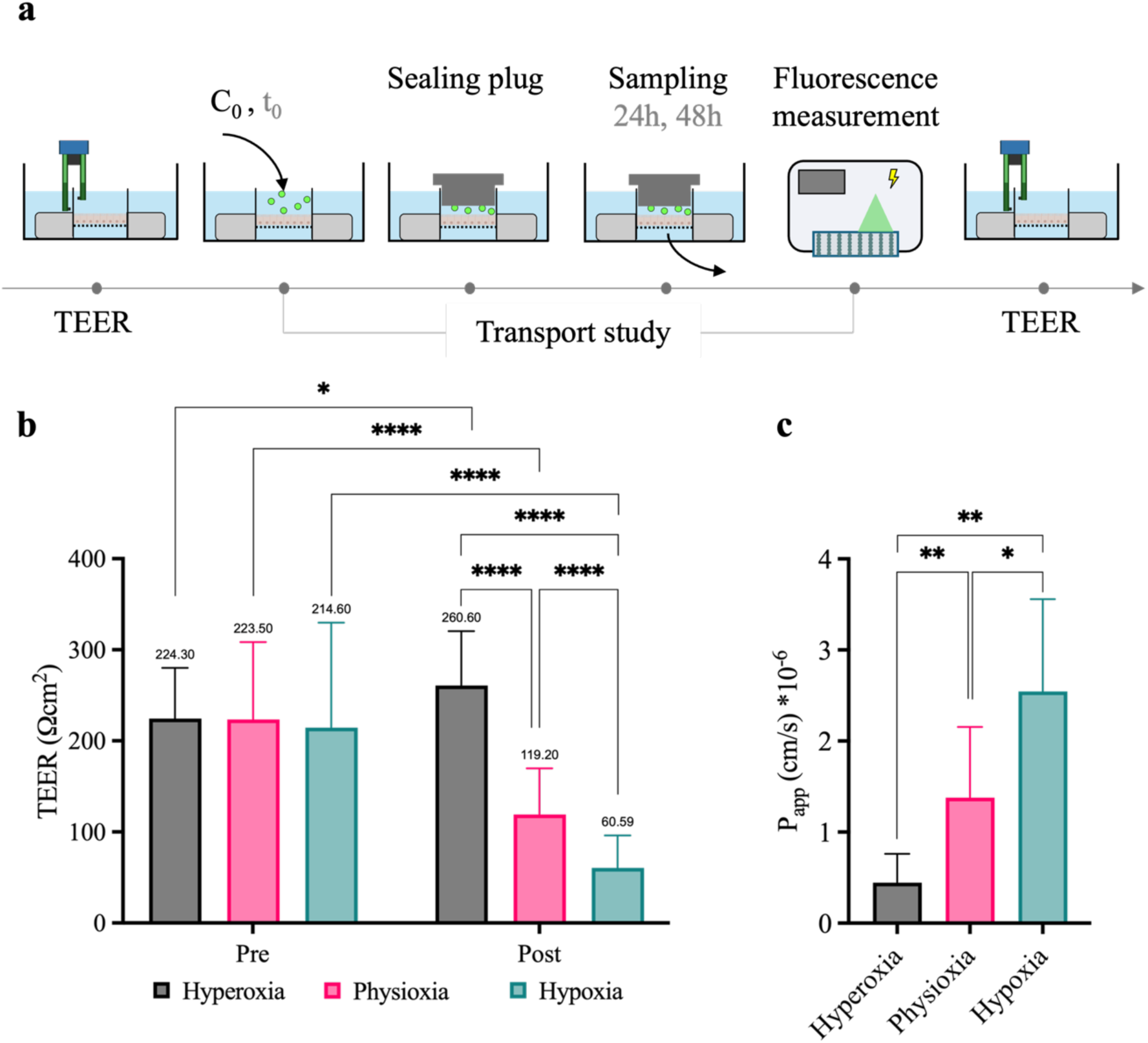
Oxygen-dependent Caco-2 cells monolayer organization. Quantitative assessment of (a) TEER (n=3 samples per condition, 3 measurements per sample), (b) cell density (n=3 samples per condition, at least 5 images per sample), and (c) tight junction length (n=3 samples per condition, at least five images per sample, 5 tight junction segment measurements per image) in different oxygen tension regimes – hyperoxia (grey), physioxia (pink) and hypoxia (teal); (d) Representative images of Caco-2 cells in the considered settings (nuclei in blue (DAPI), ZO-1 in green). Scale bar = 20 μm. Data are presented as mean ± SD. Data normality: Shapiro-Wilk test. Statistics: t-test with Welch’s correction between each pair of conditions. ** p<0.01, *** p<0.001, ****p<0.0001

## 4. Discussion

The recapitulation of physiological oxygen tension in intestinal *in vitro* models plays a pivotal role in modulating drug absorption and metabolism^[17,47]^. Although Caco-2 cells are widely recognized as a reference cellular model for drug absorption studies^[34,48,49,57,58]^, conventional Caco-2 cultures neglect the contribution of the gaseous microenvironment, thereby limiting their physiological relevance; this may have a detrimental effect on their potential in predicting *in vivo* drug transport^[48]^. At the same time, microfluidic devices and organ-on-a-chip systems are emerging as advanced *in vitro* platforms capable of more accurately recapitulating tissue-specific physiochemical cues^[59]^, including engineering strategies designed to establish physiologically relevant oxygen gradients^[18–24,60]^.

However, drug testing studies demand standardized handling procedures and adequate throughput^[34]^, requirements that are rarely fulfilled by existing microfluidic platforms incorporating controlled oxygen gradients, ultimately limiting their broad adoption in the field^[61]^.

On the other hand, the consumption of molecular oxygen by cells can be effectively exploited to induce the formation of oxygen gradients within *in vitro* models^[28–31]^, avoiding the need for bulky devices for gas exchange or complex regulation systems^[26]^, and ultimately improving devices throughput. These experimentally accessible strategies leverage the intrinsic metabolic activity of the cells themselves, positioning the biological component as the active driver of oxygen gradient formation^[47]^.

In this evolving landscape, the Gradient-on-Platform module was developed as a simple and scalable solution to induce physiologically-relevant oxygen concentrations by exploiting cellular oxygen consumption (Figure 1). The integration of the GoP module with bicompartmental or tricompartmental culture devices enabled the reproducible generation of physioxic, hyperoxic, and hypoxic conditions (Figure 2), while preserving experimental workflows and throughput comparable to standard drug absorption assays^[34,49]^.

Although some previously reported works integrating O_2_-controlled environments focused on drug metabolism^[17,60]^, the effect of physiological hypoxia on drug or other compounds (probiotics, pollutants) absorption mechanisms through the intestinal epithelium remains insufficiently evaluated. However, previous studies reported that the establishment of oxygen gradient in *in vitro* models doesn’t compromise intestinal barrier function^[29,30]^, while enabling a more faithful recapitulation of microenvironmental cues.

Here, we demonstrated that establishing a controlled oxygen gradient across intestinal barrier *in vitro* models significantly mitigated the overtight response of Caco-2 cells in mimicking drug absorption mechanisms, hence improving the *in vitro* recapitulation of compound transport for translational pre-clinical studies. Notably, this effect cannot be achieved by simply deoxygenating the culture medium or by culturing cells in an anaerobic incubator, as these approaches impose uniform hypoxic conditions^[11]^. Indeed, we further demonstrated that complete hypoxia caused a pronounced loss of intestinal barrier integrity, resulting in aberrant TEER and P_app_ values that are more representative of pathological or inflammatory states than of healthy intestinal tissue^[9,36,37,52,56]^, as an approximately 6-fold increase in P_app_ was observed relative to standard aerobic cultures, compared with the 3-fold increase measured under physioxic stimulation (Figure 4). These findings further support the need for recreating relevant compartment-specific oxygen concentrations in advanced small intestine *in vitro* models for drug development purposes^[62,63]^.

Furthermore, the establishment of a stable oxygen gradient is known to support the coculture of intestinal epithelial and endothelial cells in opposing compartments^[17,19,30]^ in microfluidic chips, therefore we hypothesize that our platform could enable the recapitulation of more complex *in vitro* models that integrate low apical oxygen tension with multicellular architectures, while maintaining compatibility with standardized Transwell-based drug absorption protocols.

## 5. Conclusion

We developed a self-sustaining approach to induce physiologically relevant oxygen concentrations in intestinal in vitro models, maintaining throughput and operating procedures comparable to standard bicompartmental inserts, for drug absorption applications. The GoP device proved to achieve target oxygen tensions for small intestine luminal and basolateral compartments, which are shown to modulate Caco-2 cells barrier function. Overall, the proposed platform represents a promising step toward the development of translational intestinal models that capture key physiological determinants of drug transport, thereby enhancing the predictive power of preclinical drug development.

## Acknowledgements

This work was supported by the National Plan for NRRP Complementary Investments (PNC, established with the decree-law 6 May 2021, n. 59, converted by law n. 101 of 2021) in the call for the funding of research initiatives for technologies and innovative trajectories in the health and care sectors (Directorial Decree n. 931 of 06-06-2022) – project n. PNC0000003 – AdvaNced Technologies for Human-centerEd Medicine (project acronym: ANTHEM). This work reflects only the authors’ views and opinions, neither the Ministry for University and Research nor the European Commission can be considered responsible for them. We sincerely appreciate the support by the National Institute for Social Security (INPS) for the PhD scholarship provided to AMAR.

## Notes

### Competing Interest Statement

Since March 2022, G.B.F. and M.S. are cofounders of TTOP Srl and hold an equity interest in the company. TTOP is developing cartridge-based technologies including the bicompartmental device used in this study. G.B.F. and M.S. are inventors on a granted Italian patent now under PCT extension (WO2022043815A1) related to the technology. The other authors declare no competing financial interests.

### Summary of Updates

Figure 2 revised: changed colors to improve readibility Minor revisions in the text to improve clarity

## References

[1] Blutt SE, Broughman JR, Zou W, Zeng XL, Karandikar UC, In J, et al. Gastrointestinal microphysiological systems. Exp Biol Med. 2017;242(16):1633–42. doi:10.1177/1535370217710638 PubMed PMID: 28534432.

[2] Zeitouni NE, Chotikatum S, von Köckritz-Blickwede M, Naim HY. The impact of hypoxia on intestinal epithelial cell functions: consequences for invasion by bacterial pathogens. Mol Cell Pediatr. 2016;3(1). doi:10.1186/s40348-016-0041-y

[3] Singhal R, Shah YM. Oxygen battle in the gut: Hypoxia and hypoxia-inducible factors in metabolic and inflammatory responses in the intestine. Journal of Biological Chemistry. 2020;295(30):10493–505. doi:10.1074/jbc.REV120.011188

[4] Ast T, Mootha VK. Oxygen and mammalian cell culture: are we repeating the experiment of Dr. Ox? Nat Metab. 2019;1(9):858–60. doi:10.1038/s42255-019-0105-0 PubMed PMID: 32694740.

[5] Keeley TP, Mann GE. Defining physiological normoxia for improved translation of cell physiology to animal models and humans. Physiol Rev. 2019;99(1):161–234. doi:10.1152/physrev.00041.2017 PubMed PMID: 30354965.

[6] Glover LE, Lee JS, Colgan SP. Oxygen metabolism and barrier regulation in the intestinal mucosa. Journal of Clinical Investigation. 2016;126(10):3680–8. doi:10.1172/JCI84429

[7] Lu Z, Imlay JA. When anaerobes encounter oxygen: mechanisms of oxygen toxicity, tolerance and defence. Nat Rev Microbiol. 2021;19(12):774–85. doi:10.1038/s41579-021-00583-y

[8] Shah YM, Matsubara T, Ito S, Yim SH, Gonzalez FJ. Intestinal Hypoxia-Inducible Transcription Factors Are Essential for Iron Absorption following Iron Deficiency. Cell Metab. 2009;9(2):152–64. doi:10.1016/j.cmet.2008.12.012

[9] Furuta GT, Turner JR, Taylor CT, Hershberg RM, Comerford K, Narravula S, et al. Hypoxia-Inducible Factor 1–Dependent Induction of Intestinal Trefoil Factor Protects Barrier Function during Hypoxia. J Exp Med. 2001;193(9):1027–34. doi:10.1084/jem.193.9.1027

[10] Corrado C, Fontana S. Hypoxia and HIF Signaling: One Axis with Divergent Effects. Int J Mol Sci. 2020;21(16):5611. doi:10.3390/ijms21165611

[11] Lian P, Braber S, Varasteh S, Wichers HJ, Folkerts G. Hypoxia and heat stress affect epithelial integrity in a Caco-2/HT-29 co-culture. Sci Rep. 2021;11(1):1–14. doi:10.1038/s41598-021-92574-5 PubMed PMID: 34162953.

[12] Burke K, Li Y. Impact of Gut Microbiota on Drug Metabolism and Absorption. Curr Pharmacol Rep. 2025;11(1). doi:10.1007/s40495-025-00429-8

[13] Dahlgren D, Lennernäs H. Intestinal Permeability and Drug Absorption: Predictive Experimental, Computational and In Vivo Approaches. Pharmaceutics. 2019;11(8):411. doi:10.3390/pharmaceutics11080411

[14] Van Breemen RB, Li Y. Caco-2 cell permeability assays to measure drug absorption. Expert Opin Drug Metab Toxicol. 2005;1(2):175–85. PubMed PMID: 16922635.

[15] Hu M, Ling J, Lin H, Chen J. Use of Caco-2 Cell Monolayers to Study Drug Absorption and Metabolism. In: Yan Z, Caldwell GW, editors. Optimization in Drug Discovery: In Vitro Methods. Totowa, NJ: Humana Press; 2004. p. 19–35. Available from: https://doi.org/10.1385/1-59259-800-5:019 doi:10.1385/1-59259-800-5:019

[16] Shah P, Jogani V, Bagchi T, Misra A. Role of Caco-2 cell monolayers in prediction of intestinal drug absorption. Biotechnol Prog. 2006;22(1):186–98. doi:10.1021/bp050208u PubMed PMID: 16454510.

[17] Lucchetti M, Aina KO, Grandmougin L, Jäger C, Pérez Escriva P, Letellier E, et al. An Organ-on-Chip Platform for Simulating Drug Metabolism Along the Gut–Liver Axis. Adv Healthc Mater. 2024;13(20). doi:10.1002/adhm.202303943 PubMed PMID: 38452399.

[18] Jing B, Wang ZA, Zhang C, Deng Q, Wei J, Luo Y, et al. Establishment and Application of Peristaltic Human Gut-Vessel Microsystem for Studying Host–Microbial Interaction. Front Bioeng Biotechnol. 2020;8(March):1–14. doi:10.3389/fbioe.2020.00272

[19] Jalili-Firoozinezhad S, Gazzaniga FS, Calamari EL, Camacho DM, Fadel CW, Bein A, et al. A complex human gut microbiome cultured in an anaerobic intestine-on-a-chip. Nat Biomed Eng. 2019;3(7):520–31. doi:10.1038/s41551-019-0397-0 PubMed PMID: 31086325.

[20] Zhang C, Yu Z, Tang H, Tian K, Jiang M, Li Z, et al. A 3D-printed multi-channel microfluidic device for precise dissolved oxygen regulation in cancer hypoxia research. Talanta. 2026;296(June 2025):128469. doi:10.1016/j.talanta.2025.128469 PubMed PMID: 40532462.

[21] Kim HJ, Huh D, Hamilton G, Ingber DE. Human gut-on-a-chip inhabited by microbial flora that experiences intestinal peristalsis-like motions and flow. Lab Chip. 2012;12(12):2165–74. doi:10.1039/c2lc40074j PubMed PMID: 22434367.

[22] Comolli J, Walsh DI, Bobrow J, Lennartz CL, Guido NJ, Thorsen T. An in vitro platform for study of the human gut microbiome under an oxygen gradient. Biomed Microdevices. 2023;25(2):14. doi:10.1007/s10544-023-00653-3

[23] Shah P, Fritz J V., Glaab E, Desai MS, Greenhalgh K, Frachet A, et al. A microfluidics-based in vitro model of the gastrointestinal human-microbe interface. Nat Commun. 2016;7(May). doi:10.1038/ncomms11535 PubMed PMID: 27168102.

[24] Shin W, Wu A, Massidda MW, Foster C, Thomas N, Lee DW, et al. A Robust Longitudinal Co-culture of Obligate Anaerobic Gut Microbiome With Human Intestinal Epithelium in an Anoxic-Oxic Interface-on-a-Chip. Front Bioeng Biotechnol. 2019;7(FEB):1–13. doi:10.3389/fbioe.2019.00013 PubMed PMID: 30792981.

[25] Shirure VS, Lam SF, Shergill B, Chu YE, Ng NR, George SC. Quantitative design strategies for fine control of oxygen in microfluidic systems. Lab Chip. 2020;20(16):3036–50. doi:10.1039/D0LC00350F

[26] Palacio-Castañeda V, Velthuijs N, Le Gac S, Verdurmen WPR. Oxygen control: the often overlooked but essential piece to create better in vitro systems. Lab Chip. 2022;22(6):1068–92. doi:10.1039/d1lc00603g PubMed PMID: 35084420.

[27] Surina S, Chmielewska A, Pratscher B, Freund P, Rodríguez-Rojas A, Burgener IA. Organ-on-a-Chip: A Roadmap for Translational Research in Human and Veterinary Medicine. Int J Mol Sci. 2025;26(21). doi:10.3390/ijms262110753 PubMed PMID: 41226787.

[28] Chen Y, Lin Y, Davis KM, Wang Q, Rnjak-Kovacina J, Li C, et al. Robust bioengineered 3D functional human intestinal epithelium. Sci Rep. 2015;5:1–11. doi:10.1038/srep13708 PubMed PMID: 26374193.

[29] Kim R, Attayek PJ, Wang Y, Furtado KL, Tamayo R, Sims CE, et al. An in vitro intestinal platform with a self-sustaining oxygen gradient to study the human gut/microbiome interface. Biofabrication. 2019;12(1):015006. doi:10.1088/1758-5090/ab446e PubMed PMID: 31519008.

[30] Grant J, Lee E, Almeida M, Kim S, LoGrande N, Goyal G, et al. Establishment of physiologically relevant oxygen gradients in microfluidic organ chips. Lab Chip. 2022;22(8):1584–93. doi:10.1039/D2LC00069E

[31] Sasaki N, Miyamoto K, Maslowski KM, Ohno H, Kanai T, Sato T. Development of a Scalable Coculture System for Gut Anaerobes and Human Colon Epithelium. Gastroenterology. 2020;159(1):388–390.e5. doi:10.1053/j.gastro.2020.03.021

[32] Coppadoro L Pietro, Lombardi M, Nicolò S, Rando AMA, Fiore GB, Foglieni C, et al. Quantitative Assessment of Gut and Vascular Barriers in TTOP, a Cartridge-Based Bicompartmental Culture Platform. Adv Mater Technol. 2026;11(1):1–14. doi:10.1002/admt.202500065

[33] Rando AMA, Russo C, Fiore GB, Soncini M. Tricompartmental platform to recapitulate epithelium-stroma-endothelium interactions. In: Convegno Nazionale di Bioingegneria. 2025.

[34] ICH. ICH Guideline ‘Biopharmaceutics Classification System-Based Biowaivers M9. M9 Guideline. 2019;(November):1–16. Available from: https://database.ich.org/sites/default/files/M9_Guideline_Step4_2019_1116.pdf

[35] Glover LE, Lee JS, Colgan SP. Oxygen metabolism and barrier regulation in the intestinal mucosa. Journal of Clinical Investigation. 2016;126(10):3680–8. doi:10.1172/JCI84429

[36] Taylor CT, Dzus AL, Colgan SP. Autocrine Regulation of Epithelial Permeability by Hypoxia: Role for Polarized Release of Tumor Necrosis Factor alpha. Gastroenterology. 1998;657–68.

[37] Friedman GB, Taylor CT, Parkos CA, Colgan SP. Epithelial permeability induced by neutrophil transmigration is potentiated by hypoxia: Role of intracellular cAMP. J Cell Physiol. 1998;176(1):76–84. doi:10.1002/(SICI)1097-4652(199807)176:1<76::AID-JCP9>3.0.CO;2-5 PubMed PMID: 9618147.

[38] Boyce MW, Kenney RM, Truong AS, Lockett MR. Quantifying oxygen in paper-based cell cultures with luminescent thin film sensors. Anal Bioanal Chem. 2016;408(11):2985–92. doi:10.1007/s00216-015-9189-x

[39] Buchwald P. FEM-based oxygen consumption and cell viability models for avascular pancreatic islets. Theor Biol Med Model. 2009;6(1). doi:10.1186/1742-4682-6-5 PubMed PMID: 19371422.

[40] Napper SA, Schubert RW. Michaelis Menten kinetics as a modeling assumption in a model of oxygen transport in heart. In: Biomedical Engineering I. Elsevier; 1982. p. 201–4. Available from: https://linkinghub.elsevier.com/retrieve/pii/B9780080288260500470 doi:10.1016/B978-0-08-028826-0.50047-0

[41] Burova I, Peticone C, De Silva Thompson D, Knowles JC, Wall I, Shipley RJ. A parameterised mathematical model to elucidate osteoblast cell growth in a phosphate-glass microcarrier culture. J Tissue Eng. 2019;10:204173141983026. doi:10.1177/2041731419830264

[42] Decleer M, Jovanovic J, Vakula A, Udovicki B, Agoua RS, Madder A, et al. Oxygen Consumption Rate Analysis of Mitochondrial Dysfunction Caused by Bacillus cereus Cereulide in Caco-2 and HepG2 Cells. Toxins (Basel). 2018;10(7):266. doi:10.3390/toxins10070266

[43] Rogers ZJ, Bhatt K, Bencherif SA. Measuring Pericellular Oxygen Tension for In Vitro Cell Culture. In: Gilkes DM, editor. Hypoxia: Methods and Protocols. New York, NY: Springer US; 2024. p. 125–31. Available from: https://doi.org/10.1007/978-1-0716-3633-6_8 doi:10.1007/978-1-0716-3633-6_8

[44] Zhang Z, Xie Y, Yi Q, Liu J, Yang L, Wang R, et al. PEAK1 maintains tight junctions in intestinal epithelial cells and resists colitis by inhibiting autophagy-mediated ZO-1 degradation. Nature Communications. 2025;16(1). doi:10.1038/s41467-025-62107-z PubMed PMID: 40707483.

[45] Kaur B, Miglioranza Scavuzzi B, F Abcouwer S, N Zacks D. A simplified protocol to induce hypoxia in a standard incubator: A focus on retinal cells. Exp Eye Res. 2023;236:109653. doi:10.1016/J.EXER.2023.109653 PubMed PMID: 37793495.

[46] Marchus CRN, Knudson JA, Morrison AE, Strawn IK, Hartman AJ, Shrestha D, et al. Low-cost, open-source cell culture chamber for regulating physiologic oxygen levels. HardwareX. 2022;11:e00253. doi:10.1016/J.OHX.2021.E00253

[47] Rinderknecht H, Ehnert S, Braun B, Histing T, Nussler AK, Linnemann C. The Art of Inducing Hypoxia. Oxygen. 2021;1(1):46–61. doi:10.3390/oxygen1010006

[48] Artursson P, Palm K, Luthman K. Caco-2 monolayers in experimental and theoretical predictions of drug transport. Adv Drug Deliv Rev. 2001;46(1–3):27–43. doi:10.1016/s0169-409x(00)00128-9 PubMed PMID: 11259831.

[49] Hubatsch I, Ragnarsson EGE, Artursson P. Determination of drug permeability and prediction of drug absorption in Caco-2 monolayers. Nat Protoc. 2007;2(9):2111–9. doi:10.1038/nprot.2007.303 PubMed PMID: 17853866.

[50] Natoli M, Leoni BD, D’Agnano I, Zucco F, Felsani A. Good Caco-2 Cell Culture Practices. Toxicology in Vitro. 2012;26(8):1243–6. doi:10.1016/j.tiv.2012.03.009 PubMed PMID: 22465559.

[51] Angelis I De, Turco L. Caco-2 Cells as a Model for Intestinal Absorption. Curr Protoc Toxicol. 2011;47(1):20.6.1–20.6.15. 10.1002/0471140856.tx2006s47

[52] Donetti E, Bendinelli P, Correnti M, Gammella E, Recalcati S, Ferraretto A. Caco2/HT-29 In Vitro Cell Co-Culture: Barrier Integrity, Permeability, and Tight Junctions’ Composition During Progressive Passages of Parental Cells. Biology (Basel). 2025;14(3). doi:10.3390/biology14030267

[53] Lopez-Escalera S, Wellejus A. Evaluation of Caco-2 and human intestinal epithelial cells as in vitro models of colonic and small intestinal integrity. Biochem Biophys Rep. 2022;31:101314. doi:10.1016/j.bbrep.2022.101314

[54] Srinivasan B, Kolli AR, Esch MB, Abaci HE, Shuler ML, Hickman JJ. TEER Measurement Techniques for In Vitro Barrier Model Systems. SLAS Technol. 2015;20(2):107–26. doi:10.1177/2211068214561025

[55] Schicho R, Krueger D, Zeller F, Von Weyhern CWH, Frieling T, Kimura H, et al. Hydrogen Sulfide Is a Novel Prosecretory Neuromodulator in the Guinea-Pig and Human Colon. Gastroenterology. 2006;131(5):1542–52. doi:10.1053/j.gastro.2006.08.035 PubMed PMID: 17101327.

[56] Nicolas A, Schavemaker F, Kosim K, Kurek D, Haarmans M, Bulst M, et al. High throughput transepithelial electrical resistance (TEER) measurements on perfused membrane-free epithelia. Lab Chip. 2021;21(9):1676–85. doi:10.1039/d0lc00770f PubMed PMID: 33861225.

[57] FDA. M9 Biopharmaceutics Classification SystemBased Biowaivers: Guidance for Industry. Published by ICH. 2021;(May):20.

[58] Artursson P, Karlsson J. Correlation between oral drug absorption and apparent drug permeability coefficients in human intetsinal epithelial (Caco-2) cells. Biochem Biophys Res Commun. 1991;175(3):880–5. Available from: http://www.sciencedirect.com/science/article/pii/0006291X9191647U

[59] Wu L, Ai Y, Xie R, Xiong J, Wang Y, Liang Q. Organoids/organs-on-a-chip: new frontiers of intestinal pathophysiological models. Lab Chip. 2023;23(5):1192–212. doi:10.1039/d2lc00804a PubMed PMID: 36644984.

[60] Lucchetti M, Kaminska M, Oluwasegun AK, Mosig AS, Wilmes P. Emulating the gut–liver axis: Dissecting the microbiome’s effect on drug metabolism using multiorgan-on-chip models. Curr Opin Endocr Metab Res. 2021;18:94–101. doi:10.1016/j.coemr.2021.03.003

[61] Malaguarnera G, Graute M, Homs Corbera A. The translational roadmap of the gut models, focusing on gut-on-chip. Open Research Europe. 2023;1:1–17. doi:10.12688/openreseurope.13709.2

[62] Li Y, Zhang H, Xiang Z, Yuan Z. Predictive Modeling of Oxygen Gradient in Gut-on-a-Chip Using Machine Learning and Finite Element Simulation. Applied Sciences. 2026;16(2):571. doi:10.3390/app16020571

[63] Wang H, Li X, Shi P, You X, Zhao G. Establishment and evaluation of on-chip intestinal barrier biosystems based on microfluidic techniques. Mater Today Bio. 2024;26(2):101079. doi:10.1016/j.mtbio.2024.101079

