## Supplementary Information for "An *in vitro* platform with self-sustaining trans-epithelial oxygen gradient to model intestinal barrier function modulation"

### Section 1 - Estimation of PLA threaded apical well oxygen diffusion coefficient ($D_{PLA}$)

To determine the effective oxygen diffusion coefficient across the device wall, transient re-oxygenation experiments were performed. The internal compartment of the PLA threaded apical well was initially equilibrated at a reduced oxygen concentration relative to ambient air, while the external environment was maintained at atmospheric conditions. Oxygen diffusion from the surrounding air into the internal chamber was continuously monitored using a 5 mm PS-PSt7-NAU oxygen sensor spot (PreSens Precision Sensing GmbH, Germany) positioned inside the chamber. The sensor was connected via an optical fiber to an OXY-4 ST (G2) oxygen meter (PreSens Precision Sensing GmbH, Germany), and measurements were recorded over time using PreSens Measurement Studio 2 software (PreSens Precision Sensing GmbH, Germany).

The concentration-time profile was then analyzed through GraphPad Prism 10 software and fitted with a one-phase association model (Figure S1):

$$C(t) = C_{\infty} - (C_{\infty} - C_0)e^{-t/\tau}$$

where $\tau$ [s] represents the characteristic time constant of the system.

Assuming predominantly one-dimensional radial diffusion across the polymer wall and minimized external mass transfer resistance in the adjacent phases, the permeability coefficient $P$ was calculated as:

$$P = \frac{VL}{A\tau}$$

Where $P$ is the oxygen permeability coefficient [$m^2$ $s^{-1}$], $V$ is the internal volume undergoing oxygenation [$m^3$], $L$ is the threaded well wall thickness [m], defined as the difference between the external and internal radii of the cylindrical structure, $A$ is the effective surface area available for radial diffusion [$m^2$], $\tau$ is the time constant obtained from the exponential fit [s].

To estimate an effective diffusion coefficient ($D_{eff}$), interfacial partition effects were not explicitly modeled and were instead incorporated into a single lumped transport parameter by assuming a unitary partition coefficient (S=1). Under this assumption, the diffusion coefficient was approximated as:

$$D_{\mathrm{eff}} = \frac{P}{S} \approx P$$

Thus, the reported value should be interpreted as an apparent diffusion coefficient describing the overall oxygen transport across the device, rather than the intrinsic molecular diffusivity of the polymer. This simplification is appropriate for device-level modeling, where the aim is to characterize global oxygen transfer behavior rather than to resolve individual diffusion and sorption contributions.

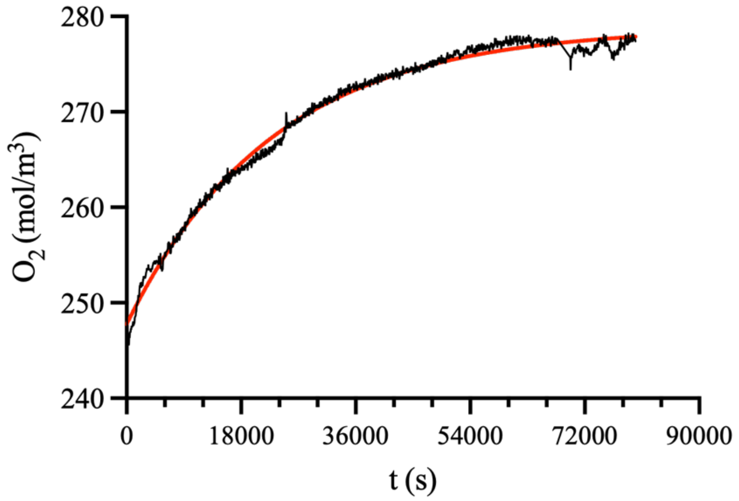


**Figure S1**. Concentration-time profile used to estimate PLA threaded apical well effective diffusion coefficient ($D_{PLA}$). The profile was fitted with a one-phase association model in GraphPad Prism 10. The red line represents the curve fitting from which τ was derived.

### Section 2 - Estimation of microporous polycarbonate (PC) membranes oxygen diffusion coefficient ($D_{Membrane}$)

The oxygen diffusion coefficient of microporous PC membranes was estimated as:

$$D_{membrane} = D_{PC}\,(1 - p) + D_{Medium}\,p$$

Where $D_{PC}$ is the oxygen diffusion coefficient of PC[1] [$m^2$ $s^{-1}$] and the $D_{medium}$ is oxygen diffusion coefficient in water-based medium[2] [$m^2$ $s^{-1}$] and $p$ is membrane porosity percentage provided by manufacturer datasheet, assuming that membrane's pores were completely filed with culture medium.

### Section 3 - Computational domains

*Physioxic conditions*

The computational model mimicking the Gradient-on-Platform was composed of

- 2 PC microporous membrane hosted in the Tri-TOP platform each represented by a cylinder of 6.5mm of diameter (corresponding to the Tri-TOP culture area diameter) and 0,022 mm of height
- 575 µm-thick cylinder of 6.5 mm diameter representing the hydrogel comprised between the membranes
- Cellular monolayer, represented by a cylinder of 6.5 mm diameter and 0,02 mm of height, estimated from mean polarized cell size in literature[3], positioned on the upper PC microporous membrane
- PLA threaded apical well represented by a hollow cylinder of 15.1mm external diameter and 6.5mm inner diameter
- Medium in the apical compartment within the threaded well, above the cell monolayer and below the sealing plug, consisting of a cylinder of 6.5 mm diameter and 2.15 mm height
- Medium in the basal compartment within the 12-well plate well where the device was placed, obtained by creating a cylinder of variable height and 22,1 mm diameter (diameter of a well of a

12-well tissue culture plate, BIOFIL) and subtracting the Tri-TOP geometry imported and the other domains

In the Gradient-on-Platform the Tri-TOP is located on a 2.2 mm PMMA C-shaped spacer to further enhance basal oxygen availability, increasing the basal medium volume (Figure S2a).

Hyperoxic and deeply hypoxic conditions were implemented by considering the same elements.

*Hyperoxic condition*

- 1 PC microporous membrane hosted in the bicompartmental TTOP platform represented by a cylinder of 6.5mm diameter (corresponding to TTOP culture area diameter) and 0,022 mm height
- Cellular monolayer, represented by a cylinder of 6.5mm diameter and 0,02 mm of height, estimated from mean polarized cell size in literature[3], positioned on the upper PC microporous membrane
- PLA threaded apical well represented by a hollow cylinder of 15.1 mm external diameter and 6.5mm inner diameter
- Medium in the apical compartment within the threaded well, above the cell monolayer and in contact with atmospheric oxygen supply
- Medium in the basal compartment within the 12-well plate well where the device was placed, obtained by creating a cylinder of variable height and 22,1 mm diameter (diameter of a well of a 12-well tissue culture plate, BIOFIL) and subtracting the bicompartmental TTOP geometry imported and the other domains

As in the Gradient-on-Platform, the bicompartmental TTOP is located on a 2.2 mm PMMA C-shaped spacer to further enhance basal oxygen availability (Figure S2b).

*Fully hypoxic condition*

- 1 PC microporous membrane hosted in the bicompartmental TTOP platform represented by a cylinder of 6.5mm diameter (corresponding to TTOP culture area diameter) and 0,022 mm height
- Cellular monolayer, represented by a cylinder of 6.5 mm diameter and 0,02 mm height, estimated from mean polarized cell size in literature[3], positioned on the upper PC microporous membrane
- PLA threaded apical well represented by a hollow cylinder of 15.1 mm external diameter and 6.5 mm inner diameter
- Medium in the apical compartment within the threaded well, above the cell monolayer and below the sealing plug, consisting of a cylinder of 6.5 mm diameter and 2.15 mm height
- Media in the basal compartment within the 12-well plate well where the device was placed, obtained by creating a cylinder of variable height and 22,1 mm diameter (diameter of a well of a 12-well tissue culture plate, BIOFIL) and subtracting the bicompartmental TTOP geometry imported and the other domains

Differently from previous conditions, the bicompartmental TTOP is positioned on the bottom of the 12-well plate to limit oxygen availability (Figure S2c)

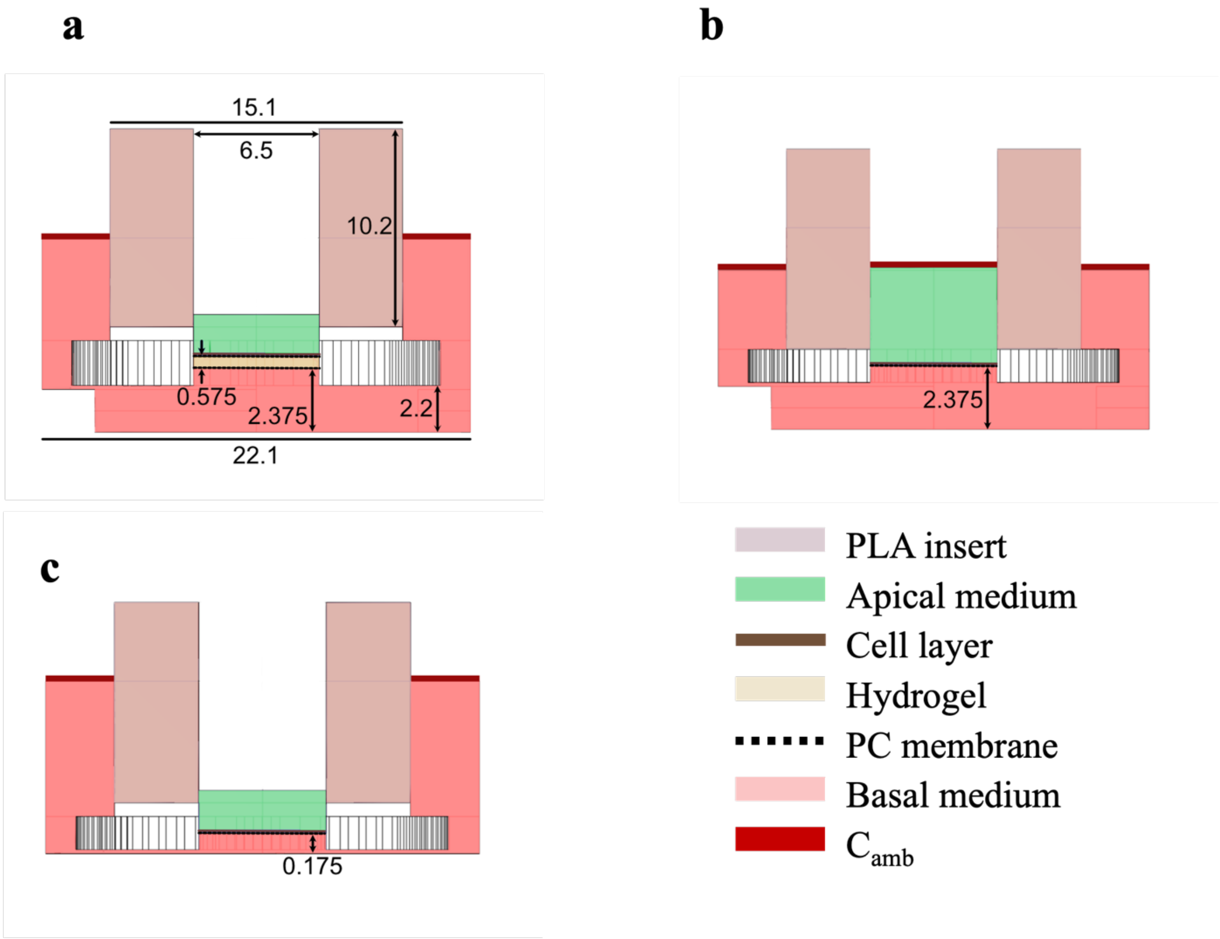


**Figure S2**. Schematic representation of the geometries imported for computational simulations of (a) physioxic, (b) hyperoxic and (c) fully hypoxic samples. All heights are reported in mm.

### References

[1] Oh JM, Begum HM, Liu YL, Ren Y, Shen K. Recapitulating Tumor Hypoxia in a Cleanroom-Free, Liquid-Pinning-Based Microfluidic Tumor Model. ACS Biomater Sci Eng. 2022;8(7):3107–21. doi:10.1021/acsbiomaterials.2c00207

[2] Buchwald P. FEM-based oxygen consumption and cell viability models for avascular pancreatic islets. Theor Biol Med Model. 2009;6(1). doi:10.1186/1742-4682-6-5 PubMed PMID: 19371422.

[3] Hidalgo IJ, Raub TJ, Borchardt RT. Characterization of the human colon carcinoma cell line (Caco-2) as a model system for intestinal epithelial permeability. Gastroenterology. 1989;96(3):736–49. PubMed PMID: 2914637
